# Strong basal/tonic TCR signals are associated with negative regulation of naive CD4^+^ T cells

**DOI:** 10.1101/2022.04.20.488956

**Authors:** Wendy M. Zinzow-Kramer, Joel Eggert, Christopher D. Scharer, Byron B. Au-Yeung

**Affiliations:** Division of Immunology, Lowance Center for Human Immunology, Department of Medicine, Emory University School of Medicine, Atlanta, GA 30322; Department of Microbiology and Immunology, Emory University School of Medicine, Atlanta, GA 30322

## Abstract

Tonic TCR signaling occurs constitutively in response to self-peptides presented by MHC (pMHC). Tonic TCR signal intensity correlates with Nur77-GFP reporter transgene expression. A broad range of Nur77-GFP is first detectable in post-selection thymocytes and persists in mature T cells. Nur77-GFP^HI^ Ly6C^−^ CD4^+^ T cells experience the strongest tonic TCR signaling and exhibit functional hypo-responsiveness to foreign pMHC stimulation. Gene expression analyses suggest similarities between the programs induced by strong tonic TCR signaling and T cell activation. However, the strongest tonic TCR signals also appear to induce expression of negative regulators, including coinhibitory receptors. Analysis of chromatin accessibility similarly suggest that strong tonic TCR signaling correlates with differentially higher accessibility of over 3000 chromatin regions in or near genes that encode positive and negative regulators of T cell activation. We propose that very strong tonic TCR signaling induces mechanisms of negative feedback to recalibrate T cell sensitivity.

## INTRODUCTION

T cells have an important role in protection from infection by recognizing foreign peptide antigens presented by MHC (pMHC). During their development, however, T cells are positively selected in the thymus for expressing TCRs that weakly recognize self-peptide antigens (self-pMHC) [1, 2]. Mature T cells retain the capacity for TCR:self-pMHC recognition in the periphery, resulting in “tonic” or “basal” TCR signaling [3]. For example, the ζ chain of the TCR complex is constitutively tyrosine phosphorylated and bound by the tyrosine kinase ZAP-70 in vivo [4, 5]. The strength of tonic TCR signaling is variable at a single cell level in naive CD4^+^ and CD8^+^ T cells. The heterogeneity of tonic TCR signal strength in naive CD4^+^ cells has been studied using surrogate markers, such as CD5, Ly6C, and Nur77-GFP [6-9]. The Nur77-GFP reporter transgene is expressed upon antigen receptor stimulation, in relative proportion to the strength of the stimulus [8, 10, 11]. It has been appreciated that the strength of tonic TCR signaling influences T cell activation and effector function [3, 12]. However, the mechanisms by which tonic signaling does so are incompletely understood.

Previously, we showed that naive CD4^+^ cells express a broad range of Nur77-GFP in the basal state, even if they express identical transgenic TCRs [13]. In line with the capacity to report on TCR signal strength, our previous studies showed that tyrosine phosphorylation of the ζ-chains was weakest in naive (CD44^LO^ Foxp3^Neg^) CD4^+^ cells with a Nur77-GFP^LO^ Ly6C^+^ phenotype, whereas ζ-chain phosphorylation was strongest in Nur77-GFP^HI^ Ly6C^−^ cells [13]. Among all naive CD4^+^ cells, the Nur77-GFP^LO^ subset mounted the most robust IL-2 and proliferative responses relative to Nur77-GFP^HI^ cells [13]. Here, we find that increasing tonic TCR signal strength correlates with gene expression and chromatin accessibility patterns suggestive of T cell activation, but also negative regulation. Together, these results suggest that the strongest tonic TCR signals are accompanied by adaptations including transcriptional and epigenetic changes that re-tune responsiveness to TCR stimuli.

## RESULTS

### A broad range of tonic TCR signal strength in the thymus and periphery

Our previous studies characterized four subpopulations of naive CD4^+^ cells (Populations A-D) based on their basal expression of Nur77-GFP and Ly6C (Fig. 1A). The frequencies of Population A (Nur77-GFP^LO^ Ly6C^+^), Population B (Nur77-GFP^MED^ Ly6C^+^), and Population C (Nur77-GFP^MED^ Ly6C^−^) cells out of the total naive CD4^+^ cell population were similar in inguinal/axial lymph nodes (LN), the red (RP) and white (WP) pulp of the spleen, and peripheral blood (PB) (Fig. 1B). However, a small but consistent decrease in the percentage of Population D (Nur77-GFP^HI^ Ly6C^−^) cells was detected in the peripheral blood relative to lymph nodes and spleen. This finding is consistent with previous studies indicating that tyrosine phosphorylation of the TCR ζ chain was decreased in T cells harvested from peripheral blood [4].

**Figure 1.**
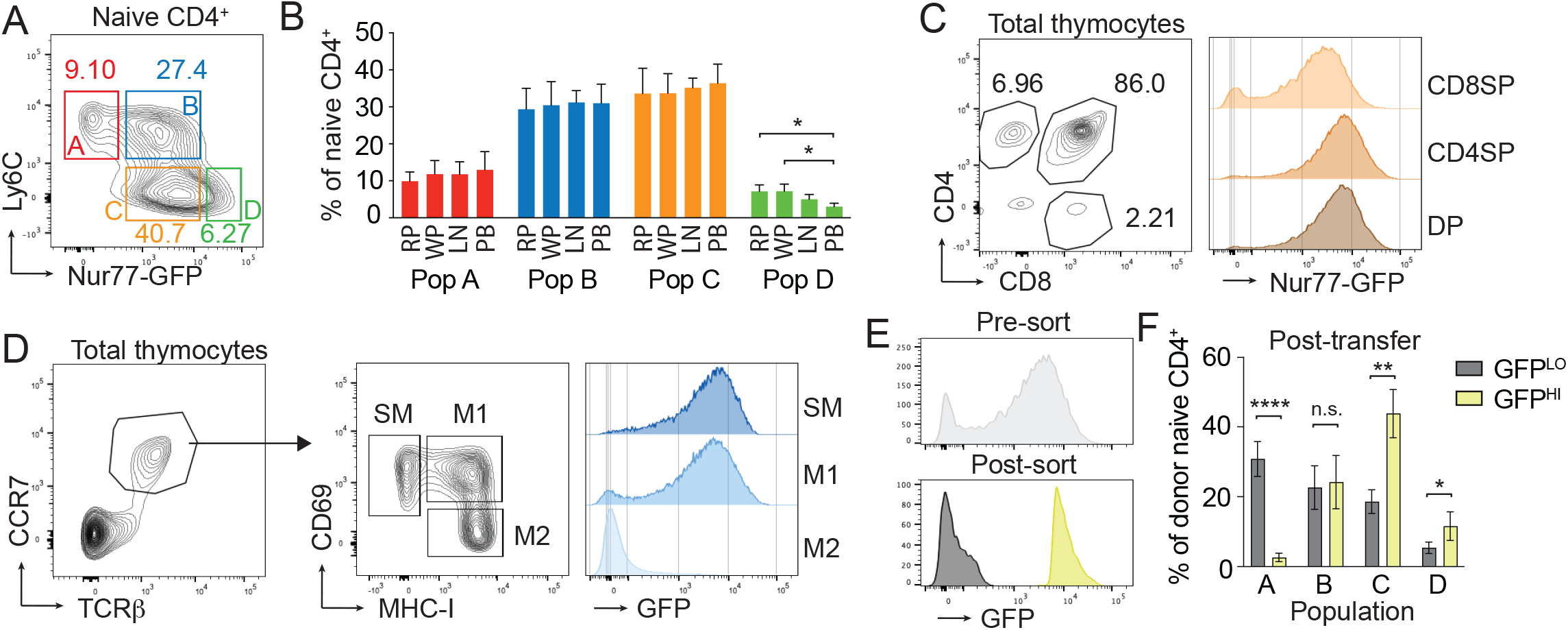
Strong tonic TCR signal strength is relatively stable. A) Contour plot shows Nur77-GFP and Ly6C expression in total naive CD4^+^ CD44^LO^ CD62L^HI^ Foxp3-RFP^neg^ cells. The four color-coded gates are labeled to indicate Populations A-D. B) Bar graph shows the percentage of Population A-D cells in the Red (RP) and White (WP) pulp of the spleen; Inguinal plus axial lymph nodes (LN); and peripheral blood (PB). Red pulp cells were identified by positive staining and white pulp cells were identified by the lack of staining following intravenous injection with anti-CD45.2 antibodies 3 minutes prior to euthanasia. C) Contour plot shows CD4 and CD8 expression by total thymocytes. Histograms show Nur77-GFP expression by the indicated populations. D) Left, Contour plot shows TCRβ and CCR7 expression by total thymocytes. Middle, Contour plot shows CD69 and MHCI expression on the TCRβ^+^ CCR7^+^ populations shown in the plot to left. Right, Histograms show Nur77-GFP expression by the indicated populations. E) Histograms show Nur77-GFP expression by total CD4SP cells (top), and purified Nur77-GFP^LO^ and Nur77-GFP^HI^ CD4SP cells (bottom). F) Bar graph shows the percentage of donor cells within the Population A-D gates 2 weeks post-adoptive transfer. N = 3 independent experiments * indicates p < 0.05; ** p<0.005; **** p<0.0001; Student’s t-Test.

We next aimed to determine whether the broad range of basal Nur77-GFP expression is also detectable in developing thymocytes. As shown previously, CD4 single-positive (CD4SP) and CD8 single-positive (CD8SP) thymocytes express higher levels of Nur77-GFP relative to double positive (DP) cells (Fig. 1C) [8, 10]. Within the CD4SP and CD8SP populations, Nur77-GFP fluorescence intensity spanned more than 2 orders of magnitude in cells, suggesting that a broad range of Nur77-GFP expression is detectable after positive selection. To compare cells at comparable stages of maturation, we analyzed thymocytes with a semi-mature (SM), mature 1 (M1), and mature 2 (M2) phenotype [14]. Each subpopulation exhibited a similarly broad spectrum of Nur77-GFP expression, suggesting that the heterogenous GFP expression level was present throughout maturation post-selection (Fig. 1D). We next asked whether relatively low or high Nur77-GFP expression in thymocytes persists in peripheral naive CD4^+^ cells. CD4SP thymocytes were sorted on the basis of low or high Nur77-GFP expression and adoptively transferred into congenic WT CD45.1^+^ hosts (Fig. 1E). Two weeks post-transfer, the majority of Nur77-GFP^LO^ donor cells were skewed toward a Population A phenotype, with graded decreases in the percentages of Population B, C, and D cells (Fig. 1F). Nur77-GFP^HI^ donor cells were skewed toward the Population C phenotype, followed by Population B and Population D (Fig. 1F). These results suggest that low or high intensity of Nur77-GFP expression in positively selected CD4^+^ cells tends to be maintained in the periphery.

### Strong basal/tonic TCR signaling is associated with decreased responsiveness to TCR stimulation

We next asked whether the differences in basal TCR signaling alter the sensitivity of Populations A-D to pMHC of varying affinities and concentrations. To address this question using the OT-II TCR transgenic system, we stimulated Population A-D cells with antigen presenting cells plus WT OVA^323-339^ peptide or an altered peptide ligand (H331R) that is less potent (Fig. 2A) [15, 16]. In response to WT OVA at relatively high concentrations, there was little detectable difference between the percentages of Population A-D cells that induced CD25 and CD69 expression (Fig. 2B and Supplemental Fig. 1A). However, stimulation with low concentrations of WT OVA or with multiple doses of H331R OVA elicited graded responses with the highest percentage of CD25^+^ CD69^+^ cells from Population A and the lowest percentage from Population D. Together these data suggest that T cells experiencing strong tonic TCR signals are relatively hypo-responsive to different doses and affinities of cognate peptide-MHC.

**Figure 2.**
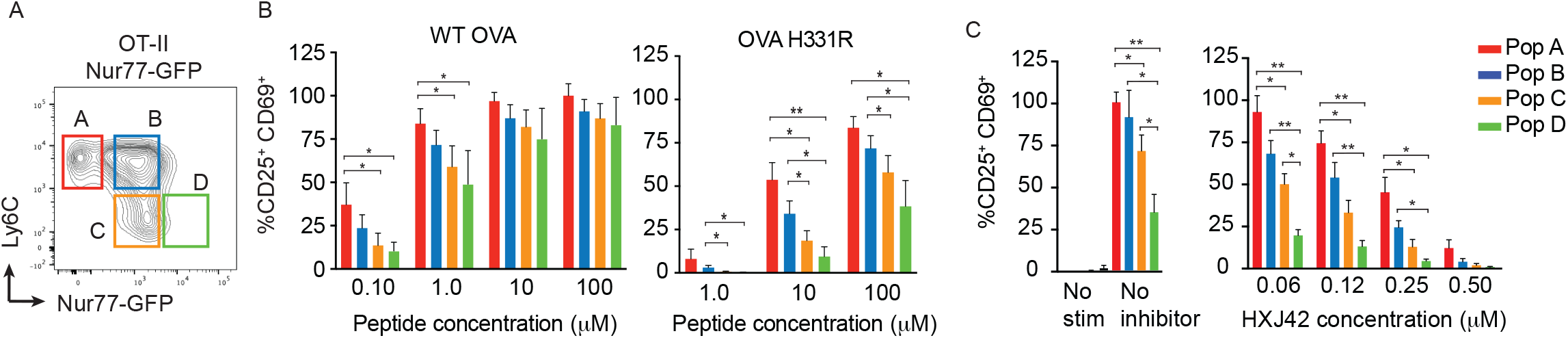
Strong tonic TCR signaling correlates with hyporesponsiveness to TCR stimulation. A) Contour plot shows Nur77-GFP and Ly6C expression by total naive CD4^+^ cells from OT-II TCR transgenic mice. Gates for Populations A-D are shown. B) Bar graphs show the percentage of CD25 and CD69 expressing cells after 24 hours of stimulation with OVA peptide or H331R peptide. C) The bar graph shows the percentage of Population A-D cells from ZAP-70^AS^ mice expressing CD25 and CD69 after 24 hours of stimulation. The concentration of HXJ42 in each condition is indicated. N = 3 independent experiments * p < 0.05; ** p<0.005 Student’s t-Test.

To assess how tonic TCR signal strength influences the sensitivity of naive T cells to inhibition of TCR-triggered signal transduction, we stimulated Population A-D cells in the presence of a ZAP-70 specific inhibitor. We took advantage of a Methionine 413 to Alanine mutant of ZAP-70 that is catalytically active but gains sensitivity to the compound HXJ42, an analog of the kinase inhibitor PP1 [17]. Purified Population A-D cells expressing the analog-sensitive ZAP-70 (ZAP-70^AS^) mutant were stimulated with the same concentration of anti-CD3 and varying concentrations of inhibitor. In the absence of inhibitor, the percentage of induced CD25 and CD69 expression progressively decreases from Population A to Population D cells (Fig. 2C and Supplemental Fig. 2B). An inhibitor concentration of 0.25 µM decreased the responses of each Population by approximately 50% although the pattern of graded decreases from the responses of Population A to Population D were still present. The concentrations of inhibitor used in these experiments did not affect CD25 and CD69 upregulation in control *Zap70*^*+/–*^ T cells, indicating that the effects of HXJ42 inhibition were specific to the analog-sensitive ZAP-70 mutant (Supplemental Fig. 2C). Together, these results suggest that as tonic TCR signaling increases, responsiveness to TCR stimuli decreases, the end result of which renders Population D cells most susceptible to inhibition of TCR signal transduction.

### Transcriptome analysis reveals upregulation of positive and negative regulators in Nur77-GFP^HI^ cells

Our results suggest that strong tonic TCR signaling correlates with attenuated responsiveness to subsequent stimulation with pMHC. To determine whether these phenotypes were associated with distinct gene expression patterns in Populations A-D, we performed bulk RNA sequencing analysis. Naive CD44^LO^ and Foxp3-negative CD4^+^ cells were sorted from non-TCR transgenic mice for this analysis. A total of 232 genes were differentially expressed (>1.5-fold) between any comparison of two populations. Principal component analysis of these differentially expressed genes suggested that Populations A and C were more similar to each other, whereas Populations B and D were most similar to each other (Fig. 3A). This pattern may reflect the common feature that Population B expresses the highest levels of Nur77-GFP amongst the Ly6C^+^ cells and Population D cells express the highest levels of Nur77-GFP amongst the Ly6C^−^ cells. The highest number of differentially expressed genes between all of the 6 pairwise comparisons occurred in the Population C vs. D comparison (Fig. 3B). We decided to focus our analysis on this comparison. Of the 174 differentially expressed genes, the majority (144) were more highly expressed in Population D, whereas 30 genes were more highly expressed in Population C (Fig. 3C). Pathway analysis of differentially expressed genes highlighted gene ontology (GO) terms that were associated with immune cell activation, proliferation, adhesion, development, and signaling (Fig. 3D). These trends suggest a gene expression profile in Population D that is associated with T cell activation. This is consistent with high Nur77-GFP expression as a surrogate marker of more intense TCR signaling in these cells, and the association between TCR signaling and the T cell activation program.

**Figure 3.**
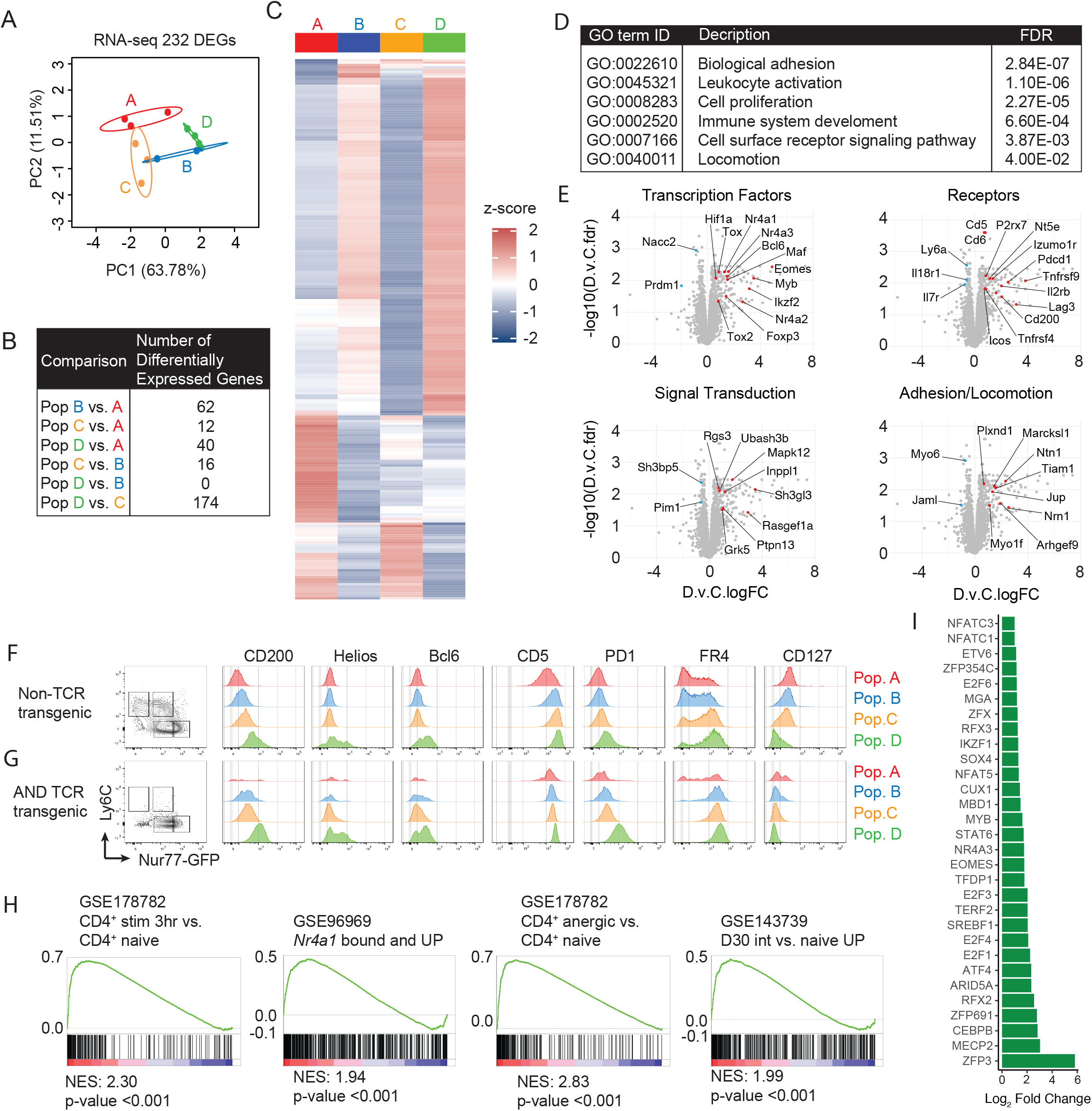
Increasing tonic TCR signal strength correlates with gene expression changes. A) Principal Component Analysis of 232 differentially expressed genes. Individual replicates corresponding to Populations A-D are indicated by separate color-coded dots. Circles denote 99% confidence intervals. B) Table shows the number of differentially expressed genes between each pairwise comparison. C) Heat map shows the relative expression levels of 232 differentially expressed genes. D) Table shows the top six gene ontology (GO) terms identified from pathway analysis of gene expression by Populations A-D. E) Volcano plots highlight subsets of differentially expressed genes within the indicated categories. F,G) Contour plots show Nur77-GFP and Ly6C expression by total naive CD4^+^ cells from TCR polyclonal and AND TCR transgenic mice. Histograms show expression of the indicated cell surface molecules for Populations A-D. H) Graphs of Gene Set Enrichment Analysis (GSEA) comparing the genes upregulated in Population D compared to Population C, versus the indicated gene sets. NES; Normalized Enrichment Score. I) Taiji analysis of transcriptional targets. Bar graph shows the transcription factors with the highest PageRank scores for the genes upregulated in Population D versus Population C.

Genes that were upregulated in Population D included *Nr4a1, Nr4a2, Nr4a3* and *Cd5*, genes that were anticipated to be upregulated in Nur77-GFP^HI^ cells. Similarly, several genes encoding co-stimulatory receptors (*Icos, Tnfrsf4, Tnfrsf9*), inhibitory receptors (*Pdcd1, Lag3, Cd200*) and markers of naturally occurring anergic T cells (*Nt5e*, encoding CD73 and *Izumo1r* encoding FR4) were upregulated in Population D. In contrast, cytokine receptor transcripts *Il7r* and *Il18r1* were decreased in Population D. Several transcription factors were upregulated in Population D, including *Bcl6, Eomes, Foxp3, Ikzf2* (Helios), *Hif1a*, and *Maf*. Genes related to cellular adhesion or cell locomotion included *Tiam, Plxnd1*, and *Arhgef9*. We verified elevated expression of several genes at the protein level on polyclonal naive CD4^+^ T cells (Fig. 3F). In addition, we took advantage of AND TCR transgenic CD4^+^ T cells, which we previously showed have skewed Nur77-GFP expression and predominantly have Population C, D phenotypes (Fig. 3G). The staining patterns of AND cells further emphasized the differential upregulation of these genes in Population D.

Gene Set Enrichment Analysis (GSEA) suggested there was a high degree of overlap between the set of genes upregulated in Population D and upregulated gene sets in naive CD4^+^ cells following acute TCR stimulation [18] or CD4^+^ cells overexpressing *Nr4a1* (Fig. 3H) [19]. This is consistent with Population D cells experiencing relatively strong TCR signaling resulting in relatively strong expression of the Nur77-GFP. Recent evidence indicates that *Nr4a1* expression is elevated in anergic and tolerized T cells [18-20]. Among the genes upregulated in Population D, there was also enrichment for gene upregulated in naturally occurring CD73^+^ FR4^+^ anergic T cells and chronically stimulated CD4^+^ cells (Fig. 3H). Analysis of the transcription factor networks upregulated in Population D revealed higher activity of Nr4a3, NFATc1 and NFATc3, transcription factors activated downstream of TCR signaling. This finding is consistent with increased TCR signal strength experienced by Population D cells (Fig. 3I). Also upregulated were targets of Eomes, which is expressed in Th1 effector cells but also expressed in dysfunctional CD4^+^ cells that have been chronically stimulated [21]. Together, these results suggest that strong tonic TCR signaling could promote gene expression changes that have features in common with activated T cells, as well as hypofunctional T cells.

### Analysis of chromatin accessibility suggests features of activation but also anergy/exhaustion/negative regulation

Considering that we observed differences at the mRNA level, we decided to investigate whether there were differences in chromatin accessibility associated with increasing tonic TCR signal strength, which might indicate epigenetic reprogramming. We performed bulk ATAC-seq analysis of Populations A-D sorted from CD4^+^ CD44^LO^ Foxp3^Neg^ cells and focused on approximately 3,200 differentially accessible regions (DAR) chromatin regions with a greater than 2-fold difference in accessibility between any two populations. Most DARs can generally be clustered into two groups: one in which accessibility is highest in Population A with a graded downward trend from Population A to Population D and a larger group of DARs with progressive increases in accessibility from Population A to Population D (Fig. 4A). The overall trends indicate a chromatin landscape that increases in accessibility as a function of increasing tonic TCR signal strength. PCA analysis of the DARs indicated that Populations A and D were most different from each other, with Populations B and C in between (Fig. 4B). Out of 3234 DARs, the vast majority, 3101, were differentially accessible between Populations A and D (Fig. 4C). Given the difference in Nur77-GFP between the Populations, we determined if changes in accessibility were associated with Nur77 binding motifs. There were 490 known Nur77 binding motifs within DARs in this dataset and the mean accessibility score increased from Population A to Population D (Fig. 4D), implying that the highest Nur77 activity occurs in Population D cells. Pathway analysis identified GO terms that were associated with cell activation (Cell activation, Biological adhesion, Cytokine production) (Fig. 4E). These results were consistent with the trends identified in RNA-seq analysis and with cells isolated based on experiencing increasing strengths of TCR signaling.

**Figure 4.**
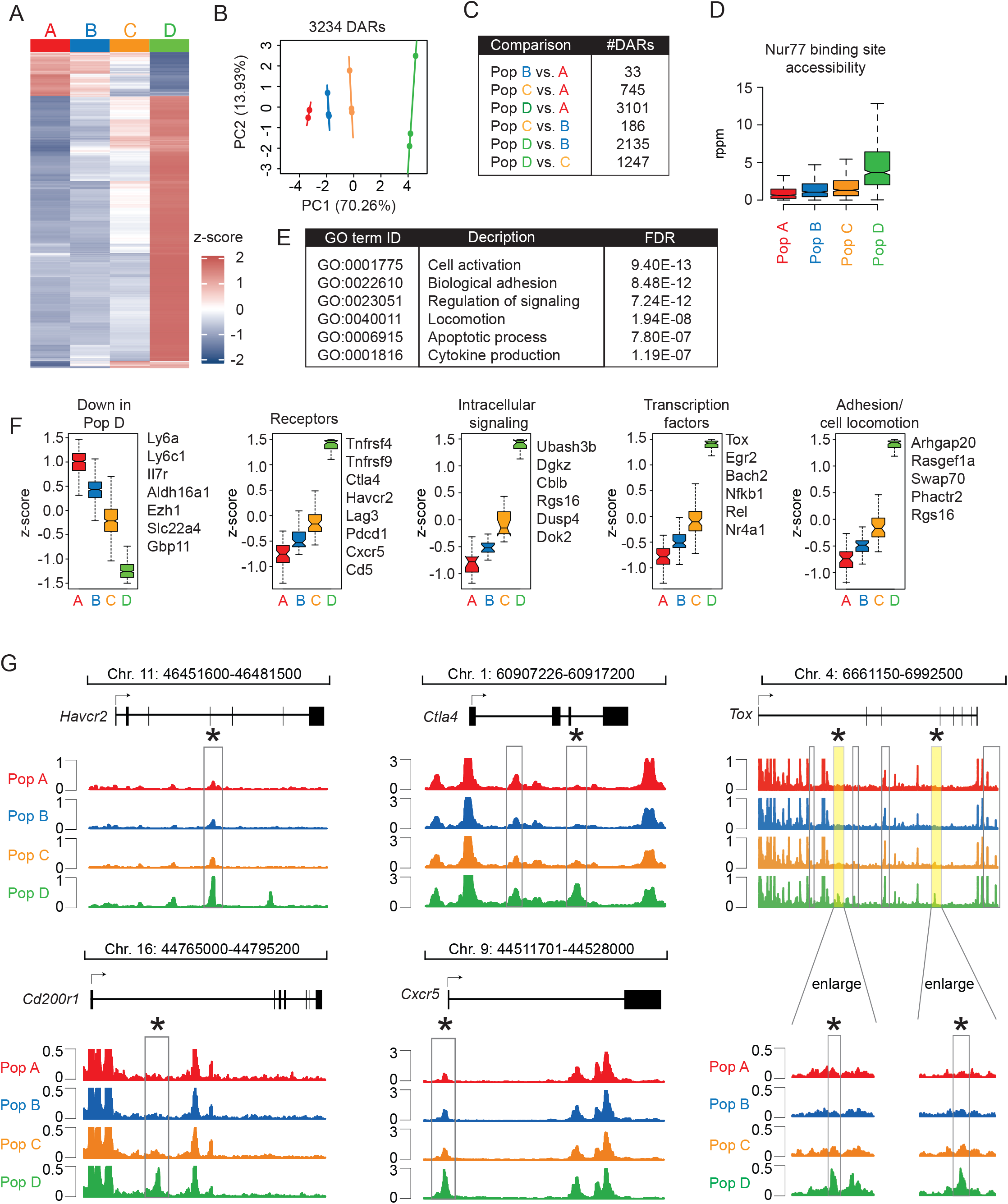
Increasing tonic TCR signal strength correlates with differences in chromatin accessibility. A) Heat map shows normalized chromatin accessibility at 3234 loci. B) PCA analysis of individual replicates from Population A-D. Circles denote 99% confidence intervals. C) Table shows the number of differentially accessible regions (DARs) between each indicated comparison A-D. D) Box and whisker graph shows the accessibility in Populations A-D of 490 Nur77 binding motifs located in DAR. F) Graphs show the average z-score values for loci located near the indicated categories of genes. To the right side of each graph is a partial list of genes with DARs in that group. G) Genome plots represent chromatin accessibility as a function of chromosome location, as determined by ATAC-seq. Each trace includes the coding region of each gene. Label at top shows the chromosome coordinates for each region. DARs are enclosed by boxes. Asterisks indicate DARs that have been previously reported to occur in effector, memory, or exhausted CD8^+^ cells. In the *Tox* locus, the DARs highlighted in yellow boxes are enlarged below to more clearly show the DARs within the selected regions.

A subset of DARs exhibited higher accessibility in Population A and lower accessibility in Population D. Among the genes that have DARs that fit this pattern included *Il7r* and *Ly6c1* (Fig. 4F). Further analysis of the *Ly6c1* locus revealed DARs located near the transcriptional start site and downstream of the coding region, both of which are least accessible in Population D (Supplemental Fig. 2A). In contrast, the majority of DARs are more highly accessible Population D relative to Populations A-C. We next analyzed DARs that are within genes encoding cell surface receptors, intracellular signaling molecules, transcription factors, and function in cell adhesion or migration. The average accessibility score increased from Population A and peaked in Population D (Fig. 4F). DARs in *Arhgap20, Rasgef1a*, and *Dgkz* were located near the first exon or first intron (Supplemental Fig. 2 B,C). Notably, several DARs were within or near genes encoding costimulatory (*Tnfrsf4* and *Tnfrsf9*) and coinhibitory receptors (*Ctla4, Havcr2, Lag3*, and *Pdcd1*) (Fig. 4F,G and Supplemental Fig. 2D). These data suggest that strong tonic signaling could promote greater accessibility of genes associated with anergy or exhaustion and have functions that are inhibitory to T cell activation.

Consistent with this correlation, we detected DARs near exon 4 of *Havcr2* (Tim3) and in introns 1 and 3 of *Ctla4*, that were more highly accessible in Population D (Fig. 4G). These DARs are similar in location to DARs that have been detected selectively in antigen-experienced cells, including effector, memory, and exhausted CD8^+^ cells [22, 23]. DARs that are detected selectively in exhausted, but not naive or effector CD8^+^ cells have been reported in intron 1 of *Cd200r1* and near exon 1 of *Cxcr5* [24]. In Population D, DARs in similar locations within *Cd200r1* and *Cxcr5* were detected (Fig. 4G). *Tox* has been implicated in CD8^+^ T cell exhaustion, and accessibility of multiple regions throughout the *Tox* locus are differentially accessible in exhausted T cells [25]. There are three DARs in intron 1, two DARs in intron 3 and several DARs outside the coding region of *Tox* in Population D. One DAR in each of intron 1 and intron 3 are in similar regions as DARs detected in exhausted CD8^+^ cells (Fig. 4G) [25]. Together, these data suggest that some features of chromatin accessibility in cells that experience strong tonic signaling are shared with chromatin accessibility in antigen-experienced T cells, including exhausted T cells.

## DISCUSSION

The strength of tonic TCR signaling can vary broadly between individual naive CD4^+^ cells. Our data support a model whereby strong tonic TCR signaling results in adaptations that include changes in gene expression and chromatin accessibility and ultimately, hypo-responsiveness to subsequent TCR stimulations.

Our data suggest that a relatively strong tone of TCR signaling in CD4SP thymocytes continues to be relatively strong after these cells localize to peripheral lymphoid organs. Nur77-GFP^HI^ cells, particularly Population D cells, may arise from incomplete negative selection of self-reactive T cells in the thymus [26-30]. We propose that chronic strong tonic TCR signaling promotes negative feedback that can lead to hyporesponsiveness or tolerance to counterbalance their increased reactivity to self-pMHC [31]. Moreover, recent studies have implicated Nr4a factors as mediators of T cell tolerance. Elevated *Nr4a1* expression is a phenotype induced by chronic TCR stimulation [21, 32] and *Nr4a* gene deficiency impairs the induction of tolerance and exhaustion [18, 19, 33, 34].

The gene expression studies shown here are in general agreement with recent work characterizing the functional and gene expression changes associated with chronic stimulation of naive CD4^+^ cells in the absence of inflammation [21]. In these studies, chronic stimulation resulted in a hypofunctional state characterized by attenuated effector cytokine potential, and a gene expression pattern associated with anergy or T cell exhaustion (*Cblb, Tox, Nr4a1, Eomes, Lag3, Nt5e, Izumo1r*, and *Pdcd1*). These genes are also upregulated in Population D, suggesting that chronic TCR stimulation in response to self-antigens may promote a gene expression program with common features.

Our analyses of chromatin accessibility highlighted DARs that are more highly accessible in Population D cells. Several of these regions, within *Ctla4, Havcr2* (Tim3), *Lag3*, and *Pdcd1* were also reported to be more highly accessible in exhausted CD8^+^ cells. Though it is currently unknown precisely which signals induce differential accessibility at these loci, we propose that the chronicity of TCR signaling experienced by both Population D cells and exhausted CD8^+^ T cells may have a role.

Among the differentially expressed genes that were upregulated in Population D were transcription factors Foxp3 and Bcl6, which are associated with Treg cells and Tfh cells, respectively. The cells that were sorted for RNA-seq analysis were sorted to exclude expression of a Foxp3-RFP reporter gene. Therefore, we conclude that Population D cells are not Treg cells but may exhibit a bias toward peripheral Treg differentiation. Previous studies have indicated that strong tonic TCR signaling, as marked by high expression of basal Nur77-GFP and CD5, or absence of Ly6C expression, correlates with a higher propensity for inducing Foxp3 expression under Treg polarizing conditions in vitro [13, 35, 36]. Similarly, the presence of elevated *Bcl6* transcript levels in Population D is suggestive of a correlation between strong tonic TCR signaling and Tfh lineage differentiation, which is consistent with recent studies [37]. These findings add support to the concept that the strength of tonic TCR signaling influences the effector differentiation of naive CD4^+^ T cells.

## ACKNOWLEDGEMENTS

We thank Julie Zikherman, Haopeng Wang, and Wan-Lin Lo for critical reading of the manuscript and Jeremy Boss for technical advice. This work was funded in part by a grant from NIH K01AR06548 (to B.A.). This study was supported in part by the Emory Flow Cytometry Core (EFCC), one of the Emory Integrated Core Facilities (EICF), and is subsidized by the Emory University School of Medicine. Additional support was provided by the National Center for Georgia Clinical & Translational Science Alliance of the National Institutes of Health under Award Number UL1TR002378.

## MATERIALS AND METHODS

### Mice

Compound mouse strains with a combination of the Nur77-GFP transgene and Foxp3-RFP knock-in allele, alone or in combination with the OT-II or AND TCR transgenes and TCRα knockout allele, have been previously described [13]. ZAP-70 M413A “analog-sensitive” (ZAP-70^AS^) mice crossed to the Nur77-GFP transgene were described previously [17]. Mice used in these studies were housed in the Division of Animal Resources at Emory University.

### Antibodies, flow cytometry analysis, and cell sorting

Thymocyte and T cell samples were analyzed using BD LSR Fortessa or FACS Symphony analyzers. Antibodies used for flow cytometry analysis were from either BD, Biolegend, or eBioscience: CD4 [clone RM4-5]; CD5 [53-7.3]; CD8α [53-6.7]; CD25 [PC61]; CD44 [IM7]; CD45.1 [A20]; CD45.2 [104]; CD69 [H1.2F3]; CD127 [SB/199]; CCR7 [4B12]; CD200 [OX-90]; PD-1 [29F.1A12]; TCRβ [H57-597]; MHCI (H-2K^b^) [AF6-88.5]; Bcl6 [K112-91]; Helios [22F6]; FR4 [eBio12A5]. Cell viability was assessed by exclusion of Live/Dead stain and intracellular staining was performed after fixation and permeabilization using the Foxp3 staining kit (Thermo Fisher). For staining of splenic red pulp cells, 3 µg anti-CD45.2 APC antibody was injected intravascularly 3 minutes prior to euthanasia, as described previously [38]. Total CD4^+^ cells were enriched by negative selection as described previously [13], followed by cell sorting with a BD FACSAria II.

### TCR stimulation

OT-II TCR transgenic cells were stimulated with T cell-depleted splenocytes and chicken ovalbumin 323-339 peptide (ISQAVHAAHAEINEAGR) or H331R peptide (ISQAVHAARAEINEAGR) (Genscript). ZAP-70 analog-sensitive T cells were stimulated with T cell depleted splenocytes and anti-CD3 antibodies [clone 145-2C11] in the presence of varying concentrations of the inhibitor HXJ42 [17].

### RNA-seq and ATAC-seq libraries

To extract RNA, 1 × 10^5^ cells were sorted directly into RLT lysis buffer (Qiagen) with 1% β-mercaptoethanol. A total of 200 pg of cDNA was used to generate a final RNA-seq library. To generate ATAC-seq libraries, a total of 2 × 10^4^ cells were sorted into media and transposed as previously described [39]. RNA-seq and ATAC-seq libraries were pooled at an equimolar ratio and sequenced at the UAB Helfin Genomics Core on a NextSeq500 using 75bp paired-end chemistry.

### RNA-seq and ATAC-seq analysis

Raw fastq files were mapped to a custom mm10 genome containing the GFP sequence using STAR [40] with the Gencode vM17 reference transcriptome. PCR duplicate reads were marked with PICARD MarkDuplicates and removed from downstream analyses. Reads mapping to exons for all unique ENTREZ genes and GFP was summarized using GenomicRanges [41] in R v3.5.2 and normalized to reads per kilobase per million. Genes expressed at 3 reads per million in all samples of one group were considered expressed. Differentially expressed genes were determined using edgeR [42], and genes that displayed an absolute log fold change > 1.5 and Benjamini-Hochberg false-discovery rate (FDR) corrected p-value < 0.05 were considered significant.

Raw ATAC-seq reads were mapped to the mm10 genome using Bowtie2 [43], enriched accessible regions for each sample determined using MACS2 [44]. A composite set of unique peaks called across all samples was annotated to the nearest gene using HOMER [45], the read depth from each sample annotated using the GenomicRanges package [41], and data normalized to reads per peak per million (rppm) in R v3.5.2. Differentially accessible regions were determined using edgeR [42]. Regions that displayed an absolute fold change > 2.0 and Benjamini-Hochberg false-discovery rate (FDR) corrected p-value < 0.05 were considered significant. Nur77 binding site accessibility was computed by first identifying the location of all Nur77 motifs using the annotatePeaks.pl [DAR file] mm10 -size given -noann -m nur77.motif - mbed nur77.motifs.bed HOMER command. Next, coverage at the 50bp surrounding each motif was calculated for all samples and the group mean for each motif summarized as a box plot using custom R scripts. Taiji analysis was performed using the standard workflow [46].

## SUPPLEMENTARY FIGURE LEGENDS

**Supplementary Figure 1**, related to Figure 2. A) Overlaid histograms show expression of Nur77-GFP, CD25, and CD69 by OT-II TCR transgenic cells sorted into Populations A-D and stimulated with the indicated concentrations of WT OVA peptide (left) or H331R peptide (right). B) Overlaid histograms show expression of Nur77-GFP, CD25, and CD69 by Populations A-D from analog-sensitive ZAP-70 mice after stimulation for 24 hours with antigen presenting cells and soluble anti-CD3 antibodies. The concentrations of the inhibitor, HXJ42, are shown on the right. C) Bar graph shows the percentage of CD25^+^ CD69^+^ cells in the presence of the indicated concentrations of HXJ42. The CD4^+^ cells were from ZAP-70 heterozygous mice (*Zap70*^*+/-*^) which express the wild-type ZAP-70 protein and serve as the analog-insensitive control cell type. **** p<0.0001; n.s. not significant p>0.05; Student’s t-Test.

**Supplementary Figure 2**, related to Figure 4. Genome trace graphs show chromatin accessibility score (y-axis) as a function of chromosomal position (x-axis). Shown are loci for the genes encoding A) Ly6C (*Ly6c1*), B) *Arhgap20* and *Rasgef1a*, C) DGKζ (*Dgkz*), and D) Lag3 in Populations A-D. Chromosome coordinates are shown below each graph.

