## Supplementary figures and images for "Strong basal/tonic TCR signals are associated with negative regulation of naive CD4^+^ T cells"

### Supplemental Figures 1-2

Supplementary Figure 1

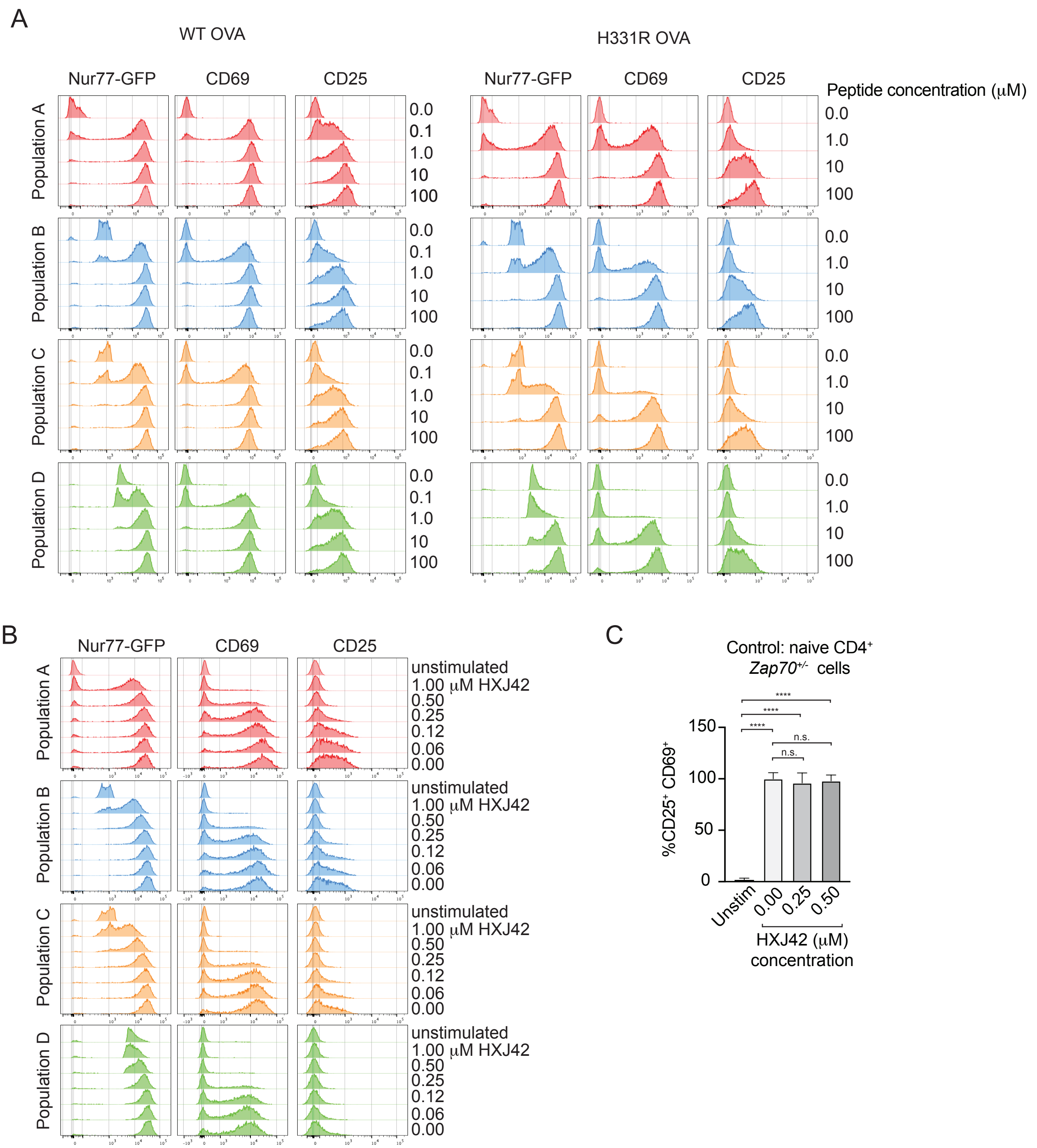

#### Supplementary Figure 2

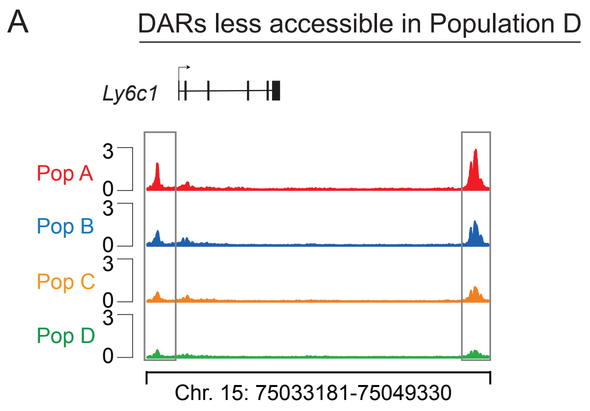


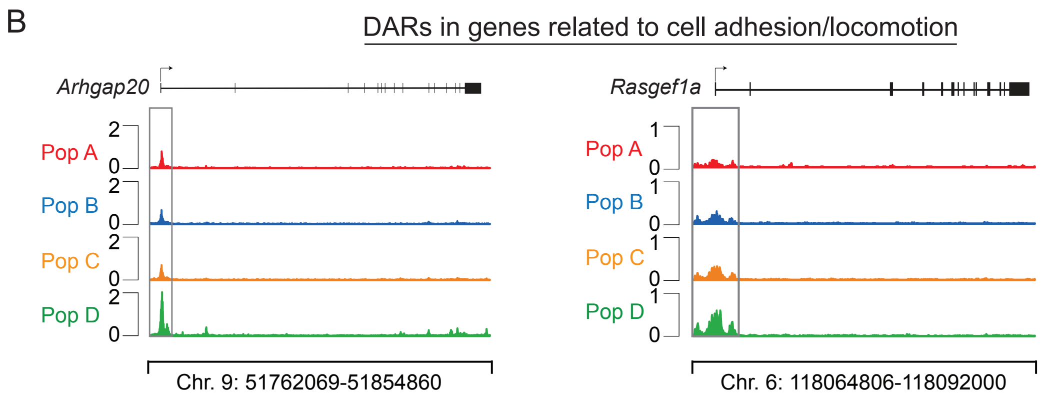


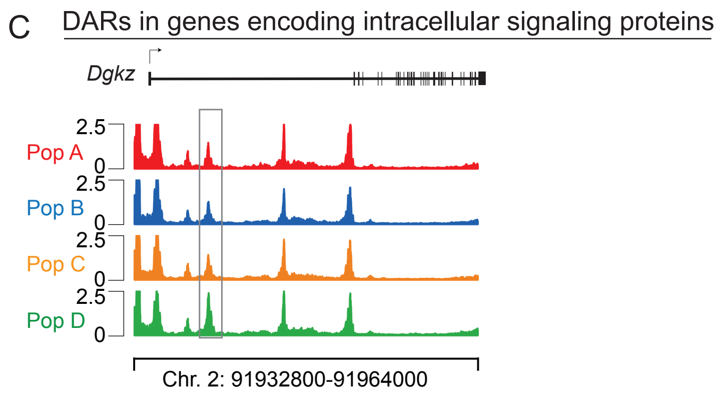


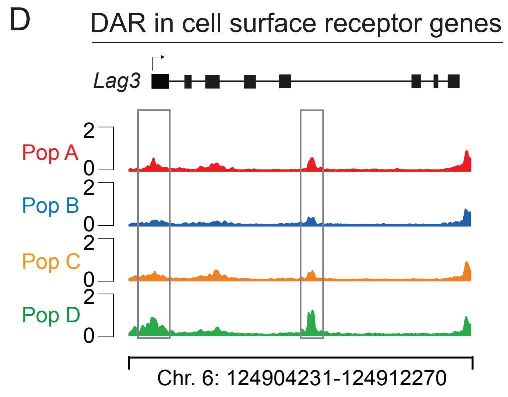
